# Whole mount multiplexed visualization of DNA, mRNA, and protein in plant-parasitic nematodes

**DOI:** 10.1101/2023.09.23.559107

**Authors:** Alexis L. Sperling, Sebastian Eves-van den Akker

## Abstract

**Background:** Plant-parasitic nematodes compromise the agriculture of a wide variety of the most common crops worldwide. Obtaining information on the fundamental biology of these organisms and how they infect the plant has been restricted by the ability to visualize intact nematodes using small molecule stains, antibodies, or *in situ* hybridization. Consequently, there is limited information available about the internal composition of the nematodes or the biology of the effector molecules they use to reprogram their host plant.

**Results:** We present the Sperling prep -a whole mount method for nematode preparation that enables staining with small molecules, antibodies, or *in situ* hybridization chain reaction. This method does not require specialized apparatus and utilizes typical laboratory equipment and materials. By dissociating the strong cuticle and interior muscle layers, we enabled entry of the small molecule stains into the tissue. After permeabilization, small molecule stains can be used to visualize the nuclei with the DNA stain DAPI and the internal structures of the digestive tract and longitudinal musculature with the filamentous actin stain phalloidin. The permeabilization even allows entry of larger antibodies, albeit with lower efficiency. Finally, this method works exceptionally well with *in situ* HCR. Using this method, we have visualized effector transcripts specific to the dorsal gland and the subventral grand of the sugar beet cyst nematode, *Heterodera schachtii*, multiplexed in the same nematode.

**Conclusion:** We were able to visualize the internal structures of the nematode as well as key effector transcripts that are used during plant infection and parasitism. Therefore, this method provides an important toolkit for studying the biology of plant-parasitic nematodes.

## Background

Plant-parasitic nematodes (PPNs) cause an annual crop loss estimated at one hundred billion American dollars worldwide [1]. Despite the huge economic losses caused by PPN infections and the threat to food security, little is known about the fundamental biology of PPNs. However, with the push towards sustainable agriculture, it is becoming increasingly important to develop new methods to control nematode populations and to do so we must understand PPN biology. PPNs have been recalcitrant to many techniques established in adjacent fields due to their biology/physiology; PPNs have a tough outer cuticle and a thick layer of muscle [2], which prevent even small molecule stains from entering the tissue.

Many of the current methods employed to visualize internal molecules or structures of PPNs require cutting the nematodes or introducing small holes into the cuticle manually in order to facilitate entry of the molecules into the tissue [3, 4]. Although perfectly adequate, there are occasions where the whole nematode is needed to assess background and specificity when determining the location of transcripts or proteins. Manually introducing small holes into the nematodes can, in some cases, also require specialized apparatus and is time consuming. There are methods that involve chemical permeabilization and electroporation of the nematodes [5, 6], which appear to work well with classical *in situ* hybridization. However, these have not proven effective with other methods of staining. In order to streamline the process of staining and *in situ* hybridization chain reactions (HCRs) in PPNs, we have developed the Sperling prep: a generalizable method of chemical and enzymatic permeabilization that allows for the entry of small molecule stains, antibodies, and HCR probes. HCR is a highly sensitive multiplexable *in situ* HCR method that works well with whole mount animals [7]. Since this method works well for a wide variety of biological visualization techniques, it simplifies the methods that need to be employed and allows for the same preparation to be used for multiple different staining procedures.

## Materials and Methods

### Nematode strain

*Heterodera schachtii* cysts were obtained from The sugar beet site of The Netherlands (IRS).

### Nematode Hatching

The second stage juvenile (J2) nematodes were hatched from isolated cysts by incubating the cysts in a 3 mM Zinc Chloride solution at 21°C. The nematodes were harvested every two days by removing the Zinc Chloride solution with the nematodes and allowing them to settle in a 50 mL tube. After the nematodes settled, the Zinc Chloride solution was decanted, and the nematodes were washed twice with 0.01% tween in water.

### Nematode Fixation

Prior to fixation the nematodes were washed 3x with M9 buffer (22 mM KH_2_PO_4_, 42 mM Na_2_HPO_4_, 20.5 mM NaCl, 1 mM MgSO_4_). The nematodes were fixed in 4% formaldehyde in M9 buffer prepared from 16% methanol-free formaldehyde (w/v), and immediately placed at -80°C for a minimum of 16 hours. Nematodes can be stored in the fixation solution at -80°C long-term. When we were ready to use them, the nematodes were removed from -80°C and allowed to defrost at 21°C for 50 min, during which the tubes were inverted 5 times by hand at 30 min and 45 min. The nematodes were centrifuged at 500 x g for 2 min and washed twice for 10 min in phosphate buffered saline (PBS) with 2% Tween20 (v/v) (2% PBST), centrifuging at 500 x g for 2 min in between.

### Cuticle Permeabilization with the Sperling preparation

The nematodes were partially dissolved in 1 ml of 1% bleach (v/v), from a 5% Sodium Hypochlorite solution, in distilled water for 2 min followed by centrifugation at 800 x g for 30 seconds. Bleach can vary in effectiveness depending on its age, storage, and light exposure; therefore, the bleach concentration and timing may need to be optimized. The bleach treatment can also be optimized to work on other species of nematodes as well as on different life stages. The nematodes were washed twice for 2 min in 2% PBST centrifuging at 500 x g for 2 min in between. As an internal control, nematodes were placed on a slide and severed with a razor using a microscope to ensure that approximately 50% of the nematodes were cut once. This was done to ensure that if the permeabilization did not work, it would nevertheless be evident whether the staining was successful. After the nematodes were severed, they were collected from the slide with 1 ml of M9 buffer and centrifuged at 500 x g for 3 min.

The collagen cuticle was then further digested with 100 μl of 10 mg/ml collagenase in 50 mM TES (10 mm Tris (pH 7.5), 10 mm Ethylenediaminetetraacetic acid (EDTA; pH 8.0), and 5% Sodium dodecyl sulfate (SDS)), supplemented with 36 mM CaCl_2_, and incubated at 37°C for 30-60 min. The tube was inverted 5 times by hand after 15 min of incubation. RNA degradation may co-occur with excessive collagenase treatment; therefore, optimization may be necessary for the timing of digestion. For small molecule staining, anywhere between 30-60 min will give adequate permeabilization. For antibody staining, a 60 min digestion will yield the best results. The HCR protocol works well when the nematodes are not cut and their RNA has not been given the opportunity to degrade, therefore 30 min is recommended for the incubation time with collagenase. After the collagenase digestion, the nematodes were first washed with 2% PBST and then washed with M9 buffer, centrifuging at 500 x g for 3 min in between. The cuticle was further digested with 5 mg/mL Proteinase K in M9 buffer at 37°C for 30 min for HCRs. A longer digestion with Proteinase K, up to 1 hour, can be used for small molecule or antibody staining. The tube was inverted 5 times by hand after 15 min of incubation. After the Proteinase K digestion, the nematodes were washed with M9 buffer and centrifuge at 500 x g for 3 min.

### Tissue Clearing and Background Quenching

After removing as much buffer as possible, the nematodes were frozen at -80°C for at least 15 min, this is a potential pause point. The nematode pellet was defrosted and 1 mL of -20°C 100% methanol was added to the tube and incubated in a -80°C ice block for 30 seconds then centrifuged for 30 seconds at 13,000 x g. The methanol was then removed and 1 mL of -20°C 100% acetone was added to the tube and incubated in a -80°C ice block for 1 min then centrifuged for 30 seconds at 13,000 x g. All but approximately 100 μL of acetone was removed and the nematodes were slowly rehydrated by adding 25 μL of distilled H_2_O every 10 min until the volume reached approximately 200 μL. Then 100 μL of 2% PBST was added to the tube and after 5 min the tube was centrifuged for 30 seconds at 4,000 x g. The nematodes were washed twice with 2% PBST and centrifuge at 500 x g for 3 min. The nematodes were then incubated in 1 mL of 2 mg/mL of glycine in distilled H_2_O for 15 min on ice. The nematodes were washed once with 2% PBST and once with 0.1% PBST, centrifuging at 500 x g for 3 min in between. At this stage the protocol splits into small molecule staining, antibody staining, or in situ HCRs.

### Small Molecule Staining

#### DNA

The nematodes were incubated in 1 μg/mL of DAPI in 200 μL 2% PBST with shaking or rotation for 16-20 hours. The excess stain was removed by washing once with 2% PBST and once 0.1% PBST for 20 min each with shaking or rotation, centrifuging at 500 x g for 3 min. Following the washes, the samples were mounted (described below).

#### Phalloidin

This staining was done after either DAPI or antibody staining but before the final wash steps. The nematodes were stained for 30 min with 5 μl of 7.3 μM phalloidin in 200 μL 2% PBST. Following the staining the nematodes were washed once with 2% PBST and once with 0.1% PBST, centrifuging at 500 x g for 3 min in between. Following the washes the samples were mounted (described below).

### Antibodies

The nematodes were blocked in 10% bovine serum albumin (BSA) (w/v) in 2% PBST for 1 hour at 21°C (room temperature) with agitation. The blocking buffer was removed and 10 μl of the primary antibody was added with 190 μl of 2% PBST and incubated 16-20 hours at 4°C with agitation. The tubes were moved to 21°C (room temperature) for 2 hours, and then moved to 37°C for 1 hour. The nematodes were washed 2 times for 20 min in 2% PBST, centrifuging at 500 x g for 3 min in between. The secondary antibody and 1 μl of DAPI were added to 200 μl 2% PBST and incubated for 16-20 hours at 4°C and 2-4 hours at 21°C (room temperature). The phalloidin staining can be done after the secondary antibody staining, if not then the nematodes proceed to the final washes. Following the staining, the nematodes were washed once with 2% PBST and once with 0.1% PBST, centrifuging at 500 x g for 3 min in between. Following the washes, the samples were mounted (described below).

Primary Antibody: (10:200 dilution) α-Mouse Myosin (5-6) from the Developmental Studies Hybridoma Bank

Secondary Antibody: (1:500 dilution) Goat α-Mouse 647 from Life Technologies.

### HCR Probes

The probes were designed by Molecular Instruments against transcript sequences we provided. For the two highly expressed targets we ordered 20 probes per transcript. Expression levels were determined from the data (in [8] Supplementary data 4).

### HCR Protocol

The method we used for *in situ* HCRs was adapted from the one provided by Molecular Instruments [7]. The Hybridization Buffer, Probe Wash Buffer, and Amplification Buffer used in the protocol were all purchased from Molecular Instruments.

Following on from the tissue clearing and background quenching, the nematodes were incubated in 50:50 Hybridization Buffer and 1% PBST at 21°C (room temperature) for 5 min. The nematodes were centrifuged at 13,000 x g for 30 seconds and the solution was carefully removed. The nematodes are found at the bottom and the top of the tube, therefore we carefully avoided aspirating them. 300 μl Hybridization Buffer was then added to the nematodes and incubated in a 37°C water bath for 1 hour. Following the prehybridization, 2 μl of the 1 μM probe stock solution and an additional 200 μl of Hybridization Buffer were added to the nematodes. The tube was then mixed well and incubated in a 37°C water bath on their side (cover cap with parafilm) for 20-22 hours. We inverted the tube 5 times by hand on two occasions during the incubation.

This portion of the method deviated from the protocol provided by Molecular Instruments because the nematodes do not easily pellet. The nematodes were at the bottom and the top of the tube, therefore we needed to avoid aspirating them. Excess probes were removed by adding 1 ml of Probe Wash Buffer that was prewarmed to 37°C and incubated for 15 min in a 37°C water bath, followed by a 13,000 x g centrifugation for 30 seconds. 1 mL of the buffer mix was removed from the tube and the process was repeated 2 more times. For the final wash with Probe Wash Buffer we removed 1.5 mL of Buffer from the tube before adding 1 ml of Probe Wash Buffer and incubating for 15 min in a 37°C water bath. The nematodes were then centrifuged at 4,000 x g for 2 min and we carefully removed all the solution. The nematodes were washed twice for 5 min with 1 mL of 5x SSCT (0.25 NaCl M, 25 mM Sodium Citrate, 0.1% Tween20) at 21°C (room temperature), centrifuged at 500 x g for 3 min in between.

The nematodes were prepared for signal amplification by incubating in 200 μL of Amplification Buffer at 21°C (room temperature) for 30 min. The amplification hairpins were prepared separately by placing 6 μL/reaction of 3 μM of the hairpin 1 (h1) stock solution in its own tube and 6 μL/reaction of 3 μM of the hairpin 2 (h2) stock solution in its own tube for each set of probe sets used. The hairpins were heated 95°C for 90 seconds and ‘Snap cooled’ to 21°C (room temperature) in a dark drawer for 30 min. The amplification hairpins were added directly to the nematodes mixing between each addition. Then an additional 100 μL of

Amplification Buffer was added to a final amplification volume of 300 μL. The nematodes were incubated for 20-22 hours in the dark at room temperature.

The excess amplification hairpins were removed by washing with 1 mL of 5x SSCT at 21°C (room temperature) twice for 5 min, then twice in 1 mL of 5x SSCT with 5 μL of DAPI for 30min, and finally once in 5x SSCT for 5 min. Following the washes, the samples were mounted.

### Mounting

Mounting the nematodes is challenging due to their size. We removed as much wash buffer as possible from nematodes in the tube. Then we removed the nematodes from the tube and placed them on a slide. A small volume of mounting media was added, roughly equivalent to 10 μL, mixed, and then the samples were secured with a cover slip and sealed with nail varnish. If too much liquid is present there will be too much space between the slide and the coverslip, which complicates imaging. Excessive liquid should be wicked off with a tissue before varnish is used to seal the coverslip in place.

### Imaging

All images were acquired on the Leica Stellaris confocal microscope with minor adjustments made to the brightness and contrast. 3D projections were created with the Leica software. The images were also prepared using ImageJ [9]. No further image manipulation was performed.

## Results

Small molecules such as DAPI or phalloidin are not able to penetrate the cuticle of fixed *H. schachtii*. If the nematodes are not physically severed, the previously developed methods do not allow small molecules to pass from the exterior to the interior of the nematode [4]. The previously developed PPN preparation methods involve fixing, cutting, enzymatic digestion, and tissue clearing. We sought to develop a method for whole mount nematodes in order to allow complete background visualization for image analysis. We therefore started with finding methods that would allow entry of DAPI, a small molecule intercalating agent used for DNA visualization, and phalloidin, a small molecule that binds to filamentous actin. We started by determining if increasing the digestion time with Proteinase K would permit stain access into the tissue. However, after 150 min the DAPI staining was limited to 33.3% (n = 30) of nematodes. Phalloidin was unable to strain the nematodes. The nematodes started to be completely degraded with increased Proteinase K digestion, therefore increasing the digestion time was deemed to not have yielded better results. We therefore added a bleach dissociation step, which is also used to remove the chorion of embryos in other species, such as *Drosophila* [10]. A 2 min 1% bleach treatment, prior to a 40 min digestion with Proteinase K increased the penetration of DAPI to 90% (n=40). Phalloidin was still unable to stain the nematodes even after that addition of the bleach treatment and the Proteinase K digestion. The cuticle is largely composed of collagen [2], hence Proteinase K digestion may not permit access past the cuticle. We therefore added a 30 min collagenase digestion in between the bleach dissolving step and the Proteinase K digestion. In order to prevent over digestion, we optimized for timing and determined that sufficient digestion could be achieved with 30 min of collagenase digestion followed by 30 min of Proteinase K digestion. This resulted in 100% (n=50) of nematodes stained with DAPI and 6% of intact nematodes stained with phalloidin. There was variation in the entry of phalloidin into the tissue. It is not clear what the basis for this variation is. Within the same preparation the digestive tract can be visualized (Figure 1a) and the longitudinal muscles can also be visualized within some nematodes (Figure 1b). When compared to the internal control of nematodes that have been severed of which 56% (n = 50) were stained strongly with phalloidin (Figure 1c-d). The staining with phalloidin was weaker in the intact nematodes but because the longitudinal muscles were not stained strongly, the staining on the internal structures of the nematodes were visible.

**Figure 1:**
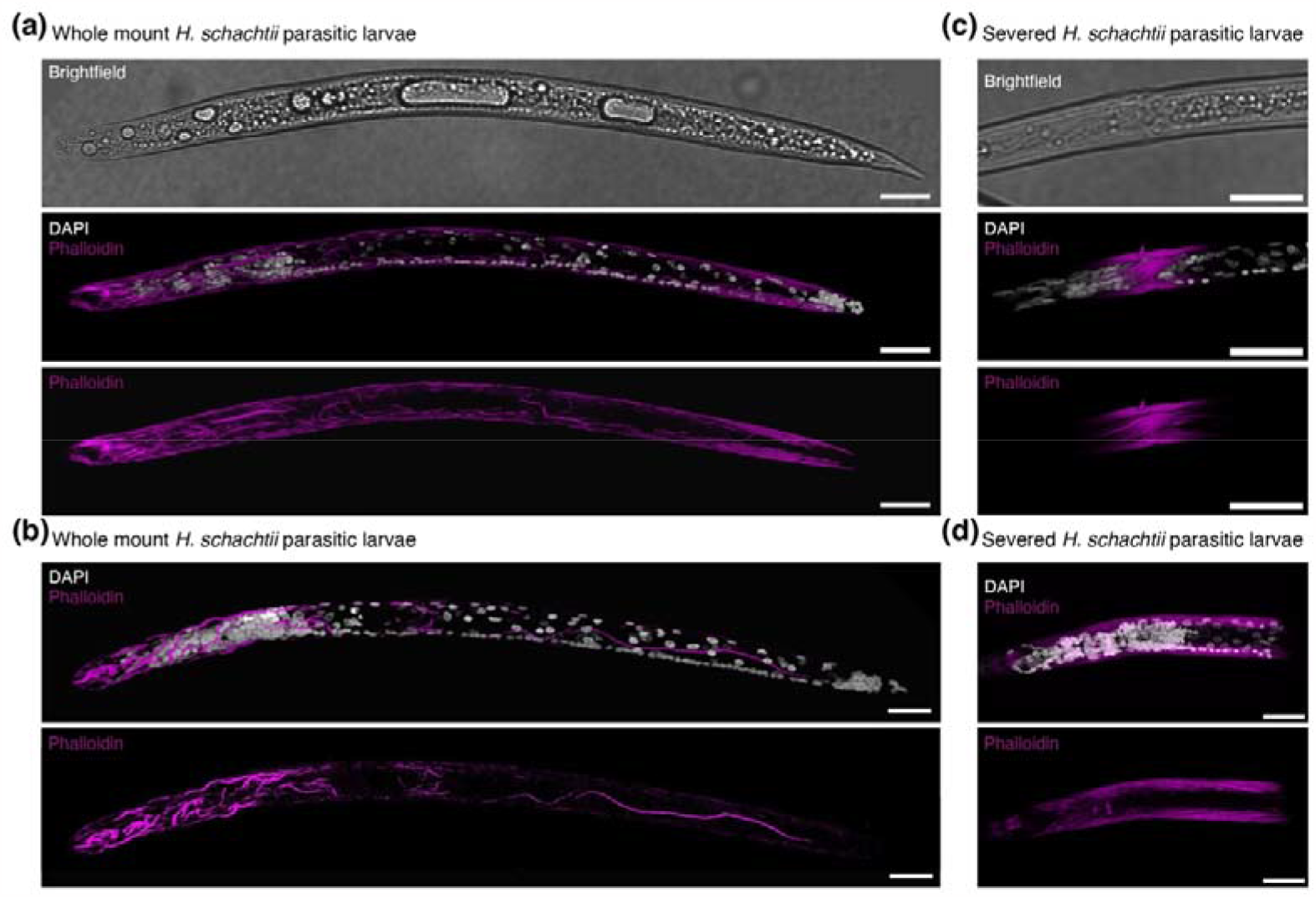
Three-dimensional projections of *H. schachtii* nematodes visualized with small molecule stains. **a-b)** Whole mount nematodes. **c-d)** Severed nematodes. The cuticle visualized with brightfield (white), DNA visualized with DAPI (white), and longitudinal muscles visualized with phalloidin (magenta). Scale bars, 20 μm.

To determine if the nematode permeabilization was sufficient to enable staining with larger molecules, we tested staining with an antibody to myosin which should bind to the longitudinal muscles of the nematodes. We found that the longitudinal muscles were stained in 52.5% of nematodes (n = 61) by extending the digestion times with collagenase and Proteinase K to 1 hour each. However, this was limited to the nematodes that were permeabilized so aggressively that their cuticle was partially opened (Figure 2a-b). 18.8% of nematodes stained with the antibody and remained largely intact (n = 32), therefore providing the whole nematode to determine the specificity of the staining. The remaining severed nematodes had much stronger antibody staining of the longitudinal muscles (Figure 2c-d).

**Figure 2:**
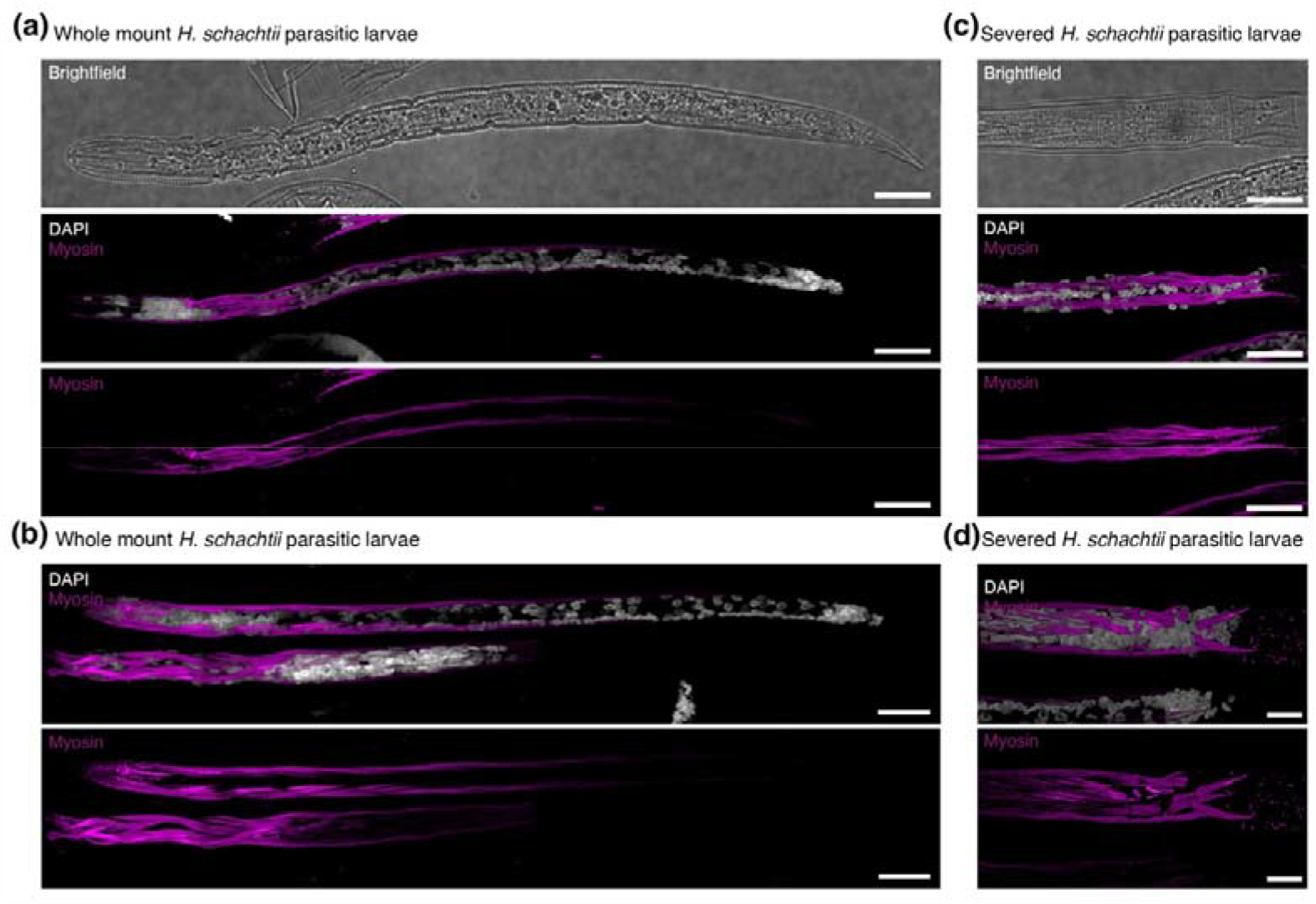
Three-dimensional projections of *H. schachtii* nematodes visualized with a small molecule and antibody stains. **a-b)** Whole mount nematodes. **c-d)** Severed nematodes. The cuticle visualized with brightfield (white), DNA visualized with DAPI (white), and longitudinal muscles visualized with an antibody generated against *C. elegans* myosin (magenta). Scale bars, 20 μm.

The ability of cyst nematodes to cause disease in plants is primarily through their “effectors” – nematode proteins primarily produced in the subventral and dorsal oesophageal gland cells that are translocated into the host plant. These effector molecules reprogram their plant host to create a syncytium that the nematode feeds from while it reproduces. There is a growing but limited understanding of what these effector molecules are; the mechanism behind the changes they induce in the plant; and how they are made. Effectors that are transported into the plant are made primarily within the two oesophageal subventral gland cells (SvG) and the one oesophageal dorsal gland cell (DG). We have developed a method to detect the expression of putative effectors within whole mount nematodes. We created an *in situ* HCR against an novel effector, Hsc_gene_2726, that is expressed in the DG of *H. schachtii*. The *in situ* HCR was present in 45.4% (n=194) of nematodes (Figure 3). A sample of the *in situ* HCRs for a single preparation shows that the expression of the effector is consistent (Figure S1). There is still variation in the strength of the staining, however, approximately half of the nematodes had sufficient hybridization with the *in situ* HCR probes to be able to determine the expression of the effector. The probe sets for *in situ* HCRs can be multiplexed, therefore we combined the DG effector with a SvG effector, the beta-1,4-endoglucanase 2 precursor *eng2*, for visualization in tandem. The *in situ* HCR for both DG effector with a SvG effector were present in 43.5% (n=216) of nematodes (Figure 4). When multiplexing, the *in situ* HCRs either worked for all probe sets or worked for none, which we attribute to a consequence of the variation in the effectiveness of the permeabilization procedure. We attribute the differences in penetration of the different visualization methods to intraindividual variation in cuticle thickness and life stage.

**Figure 3:**
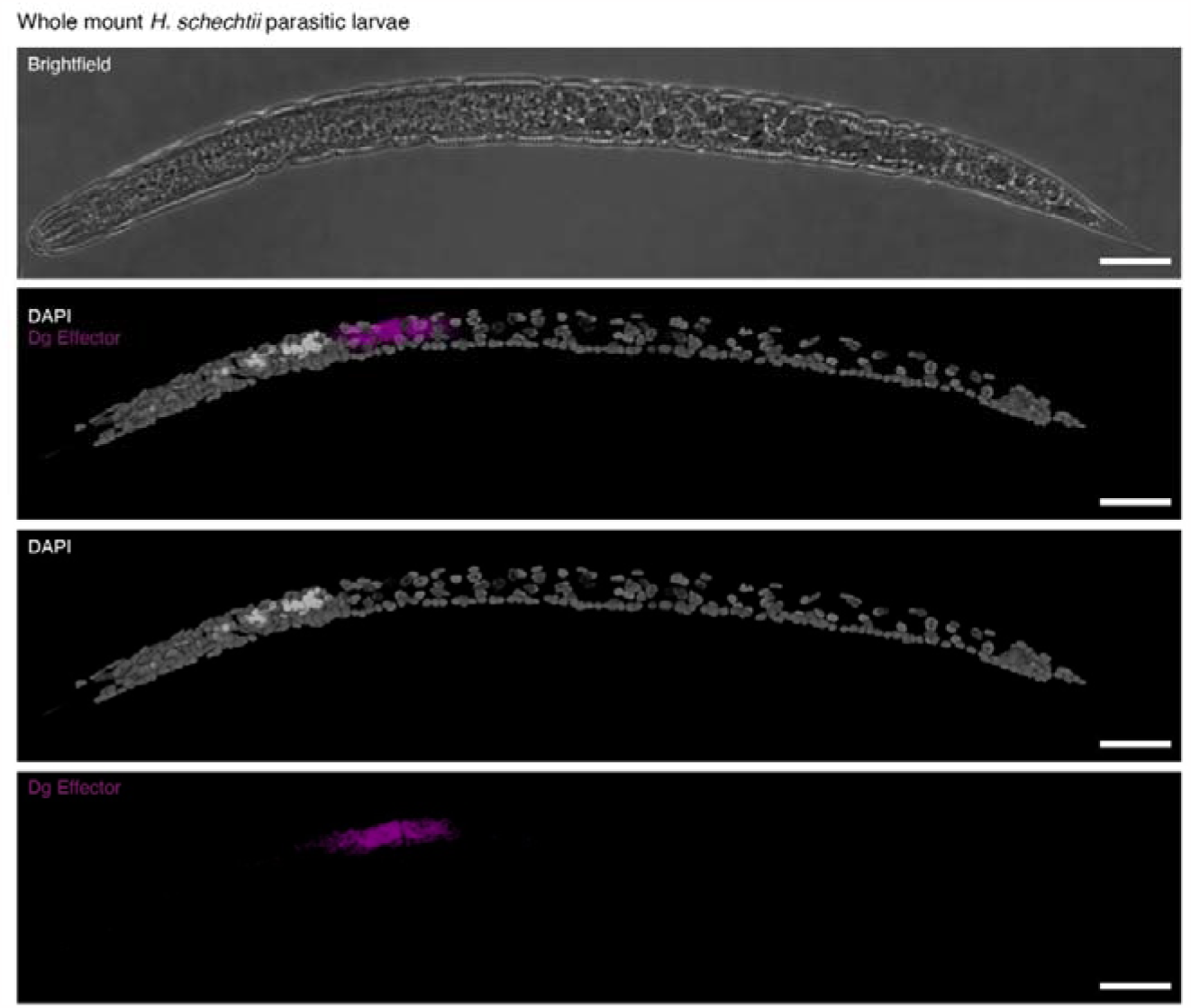
Three-dimensional projection of a whole mount *H. schachtii* nematode visualized with a small molecule stain and *in situ* HCRs. The cuticle visualized with brightfield (white), DNA visualized with DAPI (white), and DG effector Hsc_gene_2726 transcripts (magenta). Scale bars, 20 μm.

**Figure 4:**
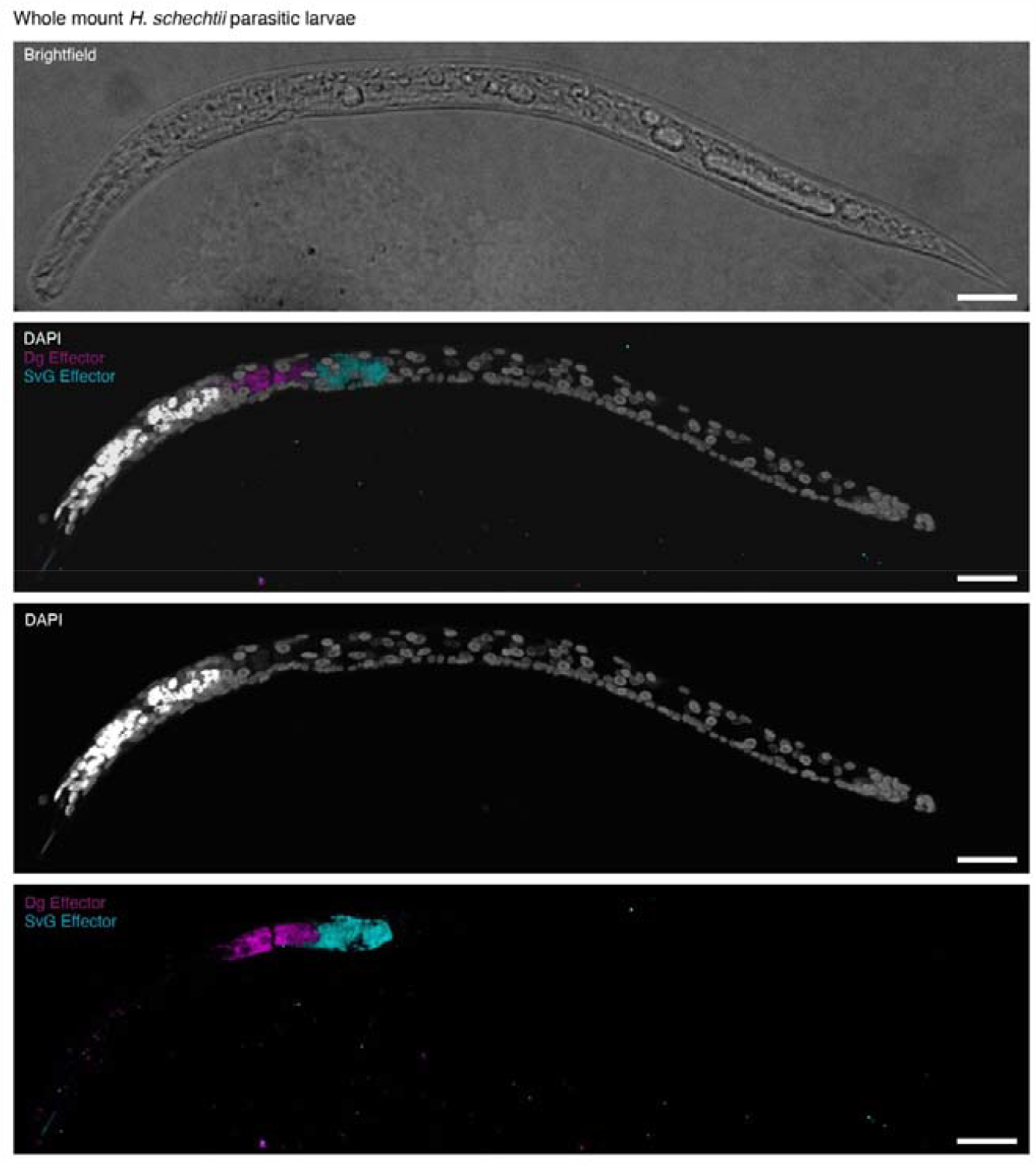
Three-dimensional projection of a whole mount *H. schachtii* nematode visualized with a small molecule stain and *in situ* HCRs. The cuticle visualized with brightfield (white), DNA visualized with DAPI (white), SvG effector Hsc_gene_21727 *eng2* transcripts (cyan), and DG effector Hsc_gene_2726 transcripts (magenta). Scale bars, 20 μm.

## Discussion

This is a highly effective and simple method to employ conventional techniques in the non-model plant-parasitic nematodes. The method that we developed requires only typical laboratory reagents and apparatus, can be performed at scale, and is highly effective. We have also simplified the process of staining and *in situ* by having the same method for preparing nematodes (Figure 5). This preparation method opens many avenues in the study of PPNs since there are many antibodies that have been generated in the model organism nematode, *Caenorhabditis elegans*, and indeed antibodies that have been generated in more distantly related species are also likely to work in PPNs if they show cross reactivity in *C. elegans*. We anticipate the introduction of *in situ* HCRs to be of the greatest interest since this method is extremely reliable and has already also been shown to work in conjunction with antibody staining in other animals. There is also very low background in the rest of the nematode (Figure 3-4), therefore it is highly specific. The size and volume of the gland cells could be visualized with the highly expressed effector transcripts (Figure 3-4). Finally, the results are unambiguous: there is no question where the genes are expressed. One of the commonly cited issues with widespread *in situ* hybridization protocols is the low success rate on a per gene basis [11] – some genes appear resistant to the protocol. In our experience, *in situ* HCR success rate is at or close to 100% and it is possible to multiplex the detection of transcripts from up to 10 with HCR at the same time in the same animal [12]. Having methods for RNA localization will be particularly important for examining the transcription of effector molecules, which are the key molecules involved in their parasitism. This method will likely work for other plant-parasitic nematodes as well as other stages of development. We have added notes within the methods sections for areas that can be optimized further to work with the species or life stage of interest.

**Figure 5:**
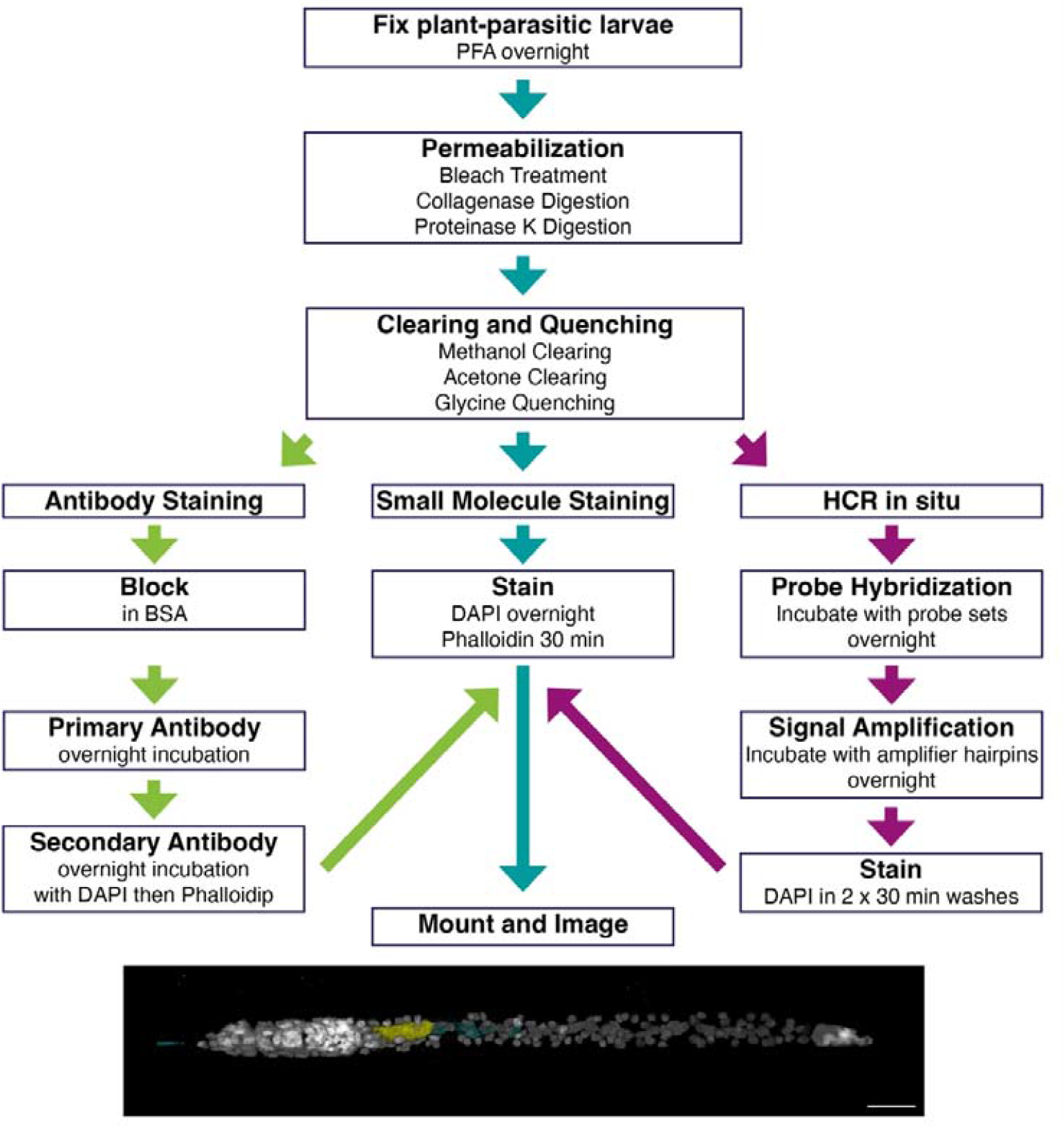
Schematic of the nematode preparation method and different staining and *in situ* HRC possibilities.

## Author contributions

ALS designed experiments, performed the experiments, collected the data, wrote the main manuscript, and analyzed the data; SEVDA contributed to the context, supervised the work, and obtained the funding. All authors read and approved the final manuscript.

## Availability of data and materials

Not applicable.

## Declarations

### Ethics approval and consent to participate

Not applicable.

## Consent for publication

Not applicable.

---

**Figure S1:**
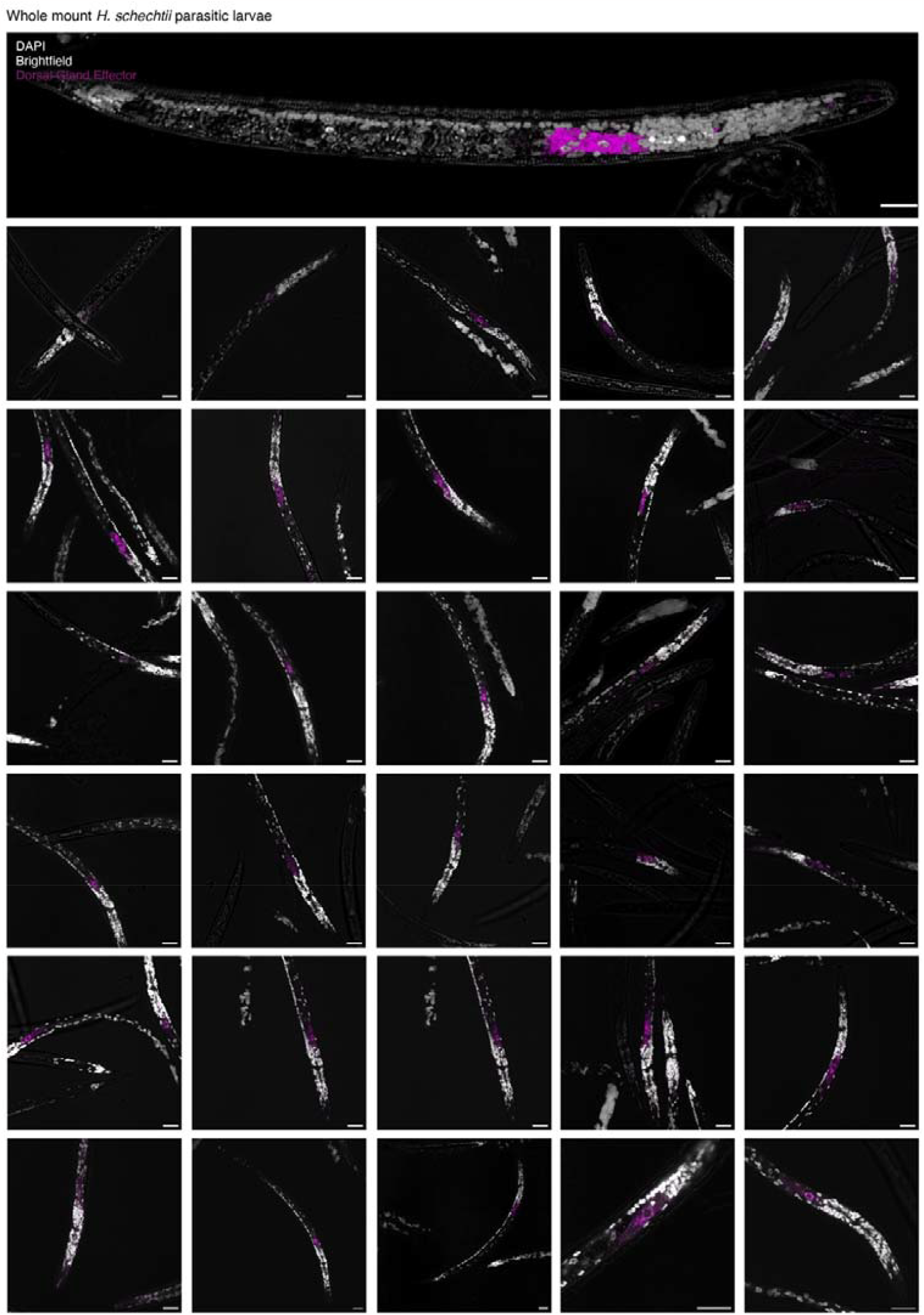
Mosaic of images of whole mount *H. schachtii* nematodes visualized with a small molecule stain and *in situ* HCRs. The cuticle visualized brightfield (white), DNA visualized with DAPI (white), and DG effector Hsc_gene_2726 transcripts (magenta). Scale bars, 20 μm.

## References

1. Nicol, J.M., et al., Current nematode threats to world agriculture. In Genomics and Molecular Genetics of Plant–Nematode Interactions (Jones, J. et al., eds) Springer, 2011: p. 21–43.

2. Basyoni, M.M. and E.M. Rizk, Nematodes ultrastructure: complex systems and processes. J Parasit Dis, 2016. 40(4): p. 1130–1140.

3. Han, Z., et al., Immobility in the sedentary plant-parasitic nematode H. glycines is associated with remodeling of neuromuscular tissue. PLoS Pathog, 2018. 14(8): p. e1007198.

4. Lilley, C.J., et al., Effector gene birth in plant parasitic nematodes: Neofunctionalization of a housekeeping glutathione synthetase gene. PLoS Genet, 2018. 14(4): p. e1007310.

5. Ruark-Seward, C.L., E.L. Davis, and T.L. Sit, Electropermeabilization-based fluorescence in situ hybridization of whole-mount plant parasitic nematode specimens. MethodsX, 2019. 6: p. 2720–2728.

6. Vandekerckhove, T.T., et al., Use of the Verrucomicrobia-specific probe EUB338-III and fluorescent in situ hybridization for detection of “Candidatus Xiphinematobacter” cells in nematode hosts. Appl Environ Microbiol, 2002. 68(6): p. 3121–5.

7. Choi, H.M., et al., Mapping a multiplexed zoo of mRNA expression. Development, 2016. 143(19): p. 3632–3637.

8. Siddique, S., et al., The genome and lifestage-specific transcriptomes of a plant-parasitic nematode and its host reveal susceptibility genes involved in trans-kingdom synthesis of vitamin B5. Nat Commun, 2022. 13(1): p. 6190.

9. Schindelin, J., et al., Fiji: an open-source platform for biological-image analysis. Nature Methods, 2012. 9(7): p. 676–682.

10. Rothwell, W.F. and W. Sullivan, Drosophila embryo dechorionation. CSH Protocols, 2007: p. pdb.prot4826.

11. de Boer, J.M., et al., In-situ Hybridization to Messenger RNA in Heterodera glycines. J Nematol., 1998. 30: p. 309–12.

12. Schulte, S.J., et al., HCR spectral imaging: 10-plex, quantitative, highresolution RNA and protein imaging in highly autofluorescent samples. bioRxiv, 2023.

